# Semi-Annual Cycles in the Biotic Communities of Temperate Aquatic Habitats

**DOI:** 10.64898/2026.05.01.721460

**Authors:** Theodor Sperlea, Conor C Glackin, Lukas Vogel, Erik Zschaubitz, Clara Nietz, Sven Karsten, Joachim W. Dippner, Stefanie Elferink, Carina Loose, Henning Schröder, Christiane Hassenrück, Matthias Labrenz

## Abstract

Recurring patterns in biosphere dynamics are anchored in daily and seasonal oscillations in abiotic variables driven by Earth’s obliquity, rotation, and orbit. While circadian and annual biotic cycles are well studied, persistent supra- or subannual cycles in biotic systems are rarely documented globally. Here, we apply a machine learning approach to DNA metabarcoding time series and detect a biotic semi-annual cycle expressed across aquatic communities in temperate regions across taxonomic domains. We propose that this dynamic reflects a semi-annual mode in insolation and is suppressed under conditions of limited nutrients or sunlight. Our results suggest photoautotrophs are central for the aetiology of the biotic SAM, while demonstrating that it is a community-level phenomena not attributable to single species. The regularity of the biotic SAM suggests value for anticipating less predictable ecological events, including phytoplankton blooms. Overall, our results highlight Earth system–scale forcing of local dynamics and reinforce coupling patterns.

## INTRODUCTION

Ecosystems are highly complex systems with intricate and sometimes chaotic dynamics (Cadotte et al. 2005, Jorgensen 2020, Levin 1998). Nevertheless, ecosystem dynamics exhibit recurrent patterns such as daily or annual cycles that can manifest at multiple levels of organization of the biotic community concurrently (Allen & Hoekstra 2015, Kamarainen & Grotzer 2019, Theise & Kafatos 2013). For example, the annual cycle affects individuals (with respond by e.g. hibernating), the population (leading to seasonal migration), and the whole community (displaying characteristically seasonal dynamics) (Kawamichi 1996, Kelly & Horton 2016, White & Hastings 2020, Wietz et al. 2021). In addition to these well-known annual cycles and across a wide range of Earth system variables, including sea levels (Zakharchuk et al. 2022), atmospheric pressure (Courtillot et al. 2022), surface and atmospheric air temperatures (Shangguan & Wang 2022, Yang & Wu 2022), and the amount of solar radi ation affecting a given area (insolation) (Paetzold & Zschörner 1961, Jones et al. 2018), the analysis of time series data has resulted in the identification of semi-annual cycles. In their ideal form, semi-annual cycles appear as sinusoidal waves with a half-year period. While some of these dynamics might emerge through interactions between intra-atmospheric processes, such as the turbulent eddies between thermospheric density variations at solstice (the “thermospheric spoon” mechanism), most are ultimately responses to patterns in insola tion (Jones et al. 2018).

Given that semi-annual modes (SAMs) are present in most abiotic variables in the Earth system, it is striking that descriptions of SAMs in biotic processes are almost completely absent from the literature. Two descriptions come in the form of independent identifications of a cycle with a six-month period in phytoplankton pigments in the offshore sea and as significant mode in a subset of phytoplankton biomass datasets, respectively (Rudjakov 1997, Winder & Cloern 2010). More recently, a biotic SAM has been described in an eight-year time series of the composition of the protist community at the SOMLIT-Astan station (Caracciolo et al. 2022). In addition to the scarcity of proof for its existence, the scope, character, and aetiology of the biotic SAM remains subject to debate. In short, whereas Rudjakov detects a SAM in a region of the earth spanning between 10°N and 10°S and links it to insolation (Rudjakov 1997), Caracciolo et al. attribute their finding to “endogenous rhythmicity or interactions between species”, describing it primarily as a local phenomenon (Caracciolo et al. 2022). Winder and Cloern, finally, note that the periodical patterns in phytoplankton can be linked to spring and autumn or summer and winter phytoplankton blooms but are “highly variable and […] may not appear regularly” (Winder & Cloern 2010). Here, we report the presence of semi-annual cycles in the dynamics of biotic communities of aquatic habitats and present a possible aetiology. Applying a machine learning approach to a one-year-long DNA metabarcoding time series sampled with high spatio-temporal resolution across freshwater, estuarine and brackish habitats, we demonstrate that the biotic SAM is not a local phenomenon but driven by semi-annual cycles in climate variables. Extending our analysis to DNA metabarcoding time series collected worldwide, we show that the biotic SAM is most likely ultimately a response to a semi-annual mode in insolation and takes effect in the temperate zone wherever the annual average concentration of Chlorophyll a surpasses a threshold. Taken together, our results show that while photoautotrophs are central to its aetiology, the biotic SAM is a community-level phenomenon.

## MATERIAL AND METHODS

### Sampling and sample analysis

Water samples were taken and analysed as described in reference 20. To control for the effect of the daily cycles, we oriented the sampling time according to sunrise, starting the sampling process 3h after sunrise. Briefly, chlorophyll a concentrations were measured using a 10-AU-005-CE fluorometer (Turner, Welschmeyer 1994) and genomic DNA was extracted from the biomass retained on a 0.22 μm Durapore PVDF filter using the KingFisher Flex purification system and the MagMAX Wastewater Kit after bead beating with MagMAX Microbiome silica beads. Paired-end DNA amplicon sequencing was performed by LGC Genomics (Berlin, Germany) on an Illumina MiSeq using the primers 5‘-CCTACGGGNGGCWGCAG-3‘ and 5‘-GACTACHVGGGTATCTAAKCC-3‘ (Herlemann et al. 2011) and 5‘-CCAGCASCYGCGGTAATTCC-3‘ and 5‘-ACTTTCGTTCTTGATYRR-3‘ (Stoeck et al. 2010) to capture the V3-V4 hypervariable region of the 16S rRNA for the Prokaryotic community and the V4 hypervariable region of the 18S rRNA gene for the Eukaryotic community, respectively. Sequence reads were further processed as documented under https://git.iow.de/bio_inf/IOWseq000042_OTCgenomics_seqs and described in reference 20. Sequences generated in the same sequencing run were used together for error learning and denoising to generate amplicon sequence variants (ASVs). For the 16S data set, which was generated from mixed orientation libraries, each orientation was denoised independently. ASVs were classified taxonomically using the RDP Bayesian classifer implemented in dada2 with the SILVA Ribosomal Reference Database version 138.1 (Quast et al. 2013) and the Protist Ribosomal Reference Database (PR2) version 4.14.0 (Guillou et al. 2013) for 16S and 18S ASVs, respectively. After combining the data from all sequencing runs into 16S and 18S ASV count tables, a rarity filter was applied to both, separately, to exclude all ASVs that were not present in at least 1% of samples with at least 0.1% of relative abundance.

### Betadiversity calculations

Bray-Curtis dissimilarities were calculated on relative ASV abundances using the vegdist function, Principal Coordinate Analyses were calculated using the betadisper function from vegan (v2.6-6) in R (v4.4.0, Oksanen et al. 2024). The amount of explained variance for each of the axes was calculated by dividing the respective eigenvalue by the sum of all eigenvalues.

### Detection of oscillations

Harmonic regression analyses were performed using the function haRmonics from the R package geoTS (v0.1.8). To assess whether a harmonic oscillation is significantly better fitting to the original data than chance, we calculated the R² between the original time series and the identified oscillation and then resampled the original time series without replacement 1000 times, performed a harmonic regression on each of the resampled time series, and calculated the R² for the resampled time series as for the original data. We finally calculated z-scores between the original harmonic’s R² and the corresponding reshuffled harmonics’ R² values. Harmonics were deemed significant if their R² had a z-score > 10, roughly corresponding to a p-value of < 10⁻⁵. To detect oscillations on the basis of the Bray-Curtis dissimilarity matrix, the pairwise dissimilarities in community composition between samples in a window of +-7 days separated by half a year were compared to all other dissimilarities using a t-test.

To detect oscillations in the multivariate datasets without a prior dimensionality reduction, machine learning models from the scikit-learn package (Pedregosa et al. 2011) version 1.2.2, in Python (v3.8) were trained to predict cosine curves generated using cos(2 * f * π * x/365 - p/180), where x is the number of days since Jan 1st of the same year, f is the frequency of the cosine oscillation, which was tested in the range 1, 2, 3, and p is the phase, which was used between 0 and 360 in steps of 360°/(data points in time series / 2). To combat the potential overestimation of model performance owing to temporal autocorrelation, we split the data points according to their collection date and used the most recent quarter of the data points to evaluate the model trained on the other three quarters. The resulting goodness-of-fit evaluation metrics are considered conservative approximations of the true statistical relationship between the oscillation in question and the total biotic community. Models were trained on the first 75% of the dataset, by date, and tested on the last 25% of the data. As metric of the performance of the model, the R² was used and calculated using the function sklearn.metrics.r2_score from the scikit-learn package. To assess significance of the model performance, null-hypothesis model performances were generated by training the same model with the same hyperparameters on cosine values reshuffled by sampling without replacement. Results were deemed significant if they exceed a z-score of 3 when compared to a distribution of 100 control points generated this way. As machine learning models, we used the ensemble.RandomForestRegressor, linear_model.Ridge and linear_model.LinearRegression from the scikit-learn package. Correlations between the relative abundance of the ASVs and the oscillations were calculated in R using the cor.test function and p-values were Bonferroni corrected.

### Insolation model

A detailed derivation of the insolation model presented in eq. 1 is in the SI. In short, assuming Earth’s orbital geometry to be circular, the amount of radiation emitted by the sun to be constant, and ignoring any atmospheric effects, we can formalize the factor I that structures the daily average insolation on a point on top of Earth’s atmosphere with colatitude or zenith angle θ at day t (where t = 0 refers to the day of the summer solstice) as I(θ, t) = 1/π Re(√(sin^2^ θ - sin^2^ δ(t)) + cos θ sin δ(t) (arcsin (tan δ(t) / tan θ) + π/2)), where sin δ(t) = sin θ_0_ cos(Ω*t*) is the angle of declination, θ_0_≈23.44° is the obliquity of Earth, and Ω = 2π/365 day⁻¹ is the annual orbiting frequency. Model output was compared to downward solar radiation flux at the nominal top of atmosphere (dswrf.ntat) data for the year 2005 from NCEP/DOE Reanalysis II. NCEP/DOE Reanalysis II data provided by the NOAA PSL, Boulder, Colorado, USA, from their website at https://psl.noaa.gov. NetCDF data was handled using the ncdf4 (v1.22) package in R.

### Additional software packages used

Figures were created using the ggplot2 (v3.5.1), reshape2 (v1.4.4), ggrepel (v0.9.5), ggpmisc (v0.5.6) and patchwork (v1.2.0) packages in R 4.4.0. Maps were generated additionally using the packages sf (v1,0-16), rnaturalearth (v1.0.1), ggspatial (v1.1.9). Furthermore, the packages data.table (v1.15.4) and plyr (v1.8.9) were used for data handling.

## RESULTS

### External factors drive the semi-annual cycle in the biotic community in the Warnow Estuary and Baltic Sea

To investigate whether the biotic SAM is a local phenomenon owing to endogenous factors, or due to external factors and thus present across multiple biotopes, we sampled surface water at fourteen sites along the Warnow River, estuary, and Baltic Sea coast twice weekly between April 2022 and April 2023 (Fig. 1C, Sperlea et al. 2025). The sampling area was chosen as it harbors a steep salinity gradient (between 0 and 23) and distinct freshwater and Baltic Sea communities that are affected by the same weather conditions but separated by a dam that prevents backflow of estuarine water into the river. We then used taxonomically broad 16S and 18S rRNA gene primers, targeting Bacteria and Eukaryota, respectively, to analyze the dynamics of the biotic community composition in surface water samples via eDNA metabarcoding (see Methods for details).

**Fig. 1:**
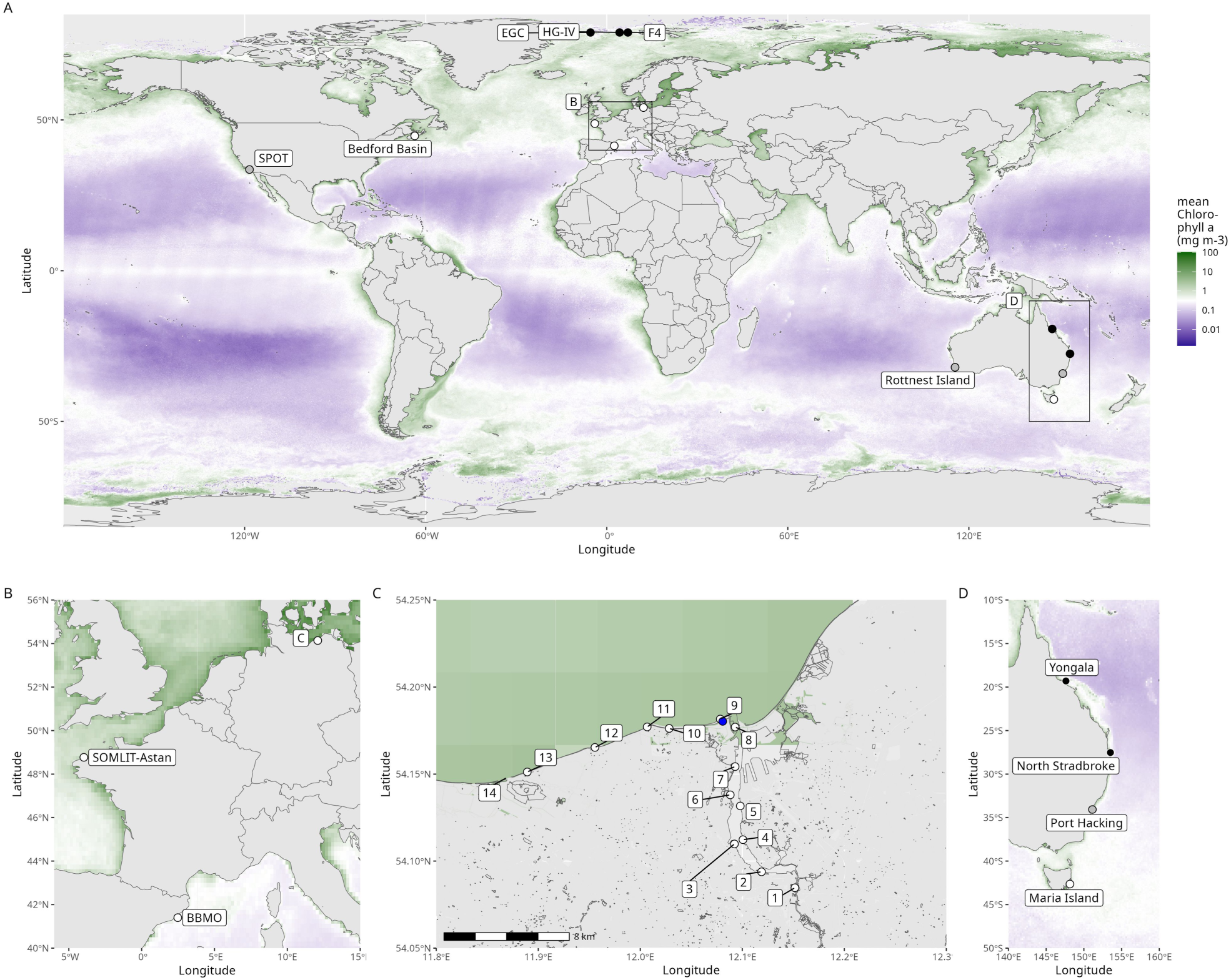
Global distribution of the time series used in this study and the annual average Chlorophyll a concentration. Dots indicate the sampling sites of the time series analyzed in this study; squares with letters indicate the regions displayed in more detail in **B**, **C**, and **D**. White dots indicate the presence, black dots the absence of a biotic SAM; grey dots indicate presence of the biotic SAM only at specific water depths. The labeled blue dot in **B** indicates the location of the DWD weather station. Colors indicate the average Chlorophyll a concentration in 2021 as given in ref. 43.

Visualizing community composition dynamics separately at each sampling site after a Bray-Curtis dissimilarity-based Principal Coordinate Analysis (PCoA) ordination, we identified patterns that resemble sinusoidal oscillations over time for all sampling sites and for both Prokaryotes and Eukaryotes among the first five PCo axes (Fig. 2). More specifically, whereas the first PCo axes always resembled annual oscillations, other PCo axes resembled semi-annual cycles, i.e., sinusoidal oscillations with two maxima and minima during the year (highlighted in Fig. 2). To confirm the presence of annually and semi-annually cyclical patterns in our data, we fit a harmonic regression model using ordinary least squares to the first five PCo axes for each sampling site and the Prokaryotic and Eukaryotic communitiesseparately. Except for sampling site 9 for 16S and site 10 for 18S, we identified at least one PCo axis that contains a semi-annual but no annual cycle at each sampling site and for both taxonomic groups (Fig. S1). As PCo axes capture orthogonal signals, this result suggests that the cycles of different frequencies are linked to different dynamics within the biotic community and potentially have distinct aetiologies. Furthermore, we were unable to detect the biotic SAM directly in the Bray-Curtis distance matrices, i.e., samples separated by half a year were not, on average, more similar than samples separated by any other period (Fig. S2). This suggests that the decomposition of community dynamics into orthogonal components is necessary to identify the biotic SAM.

**Fig. 2:**
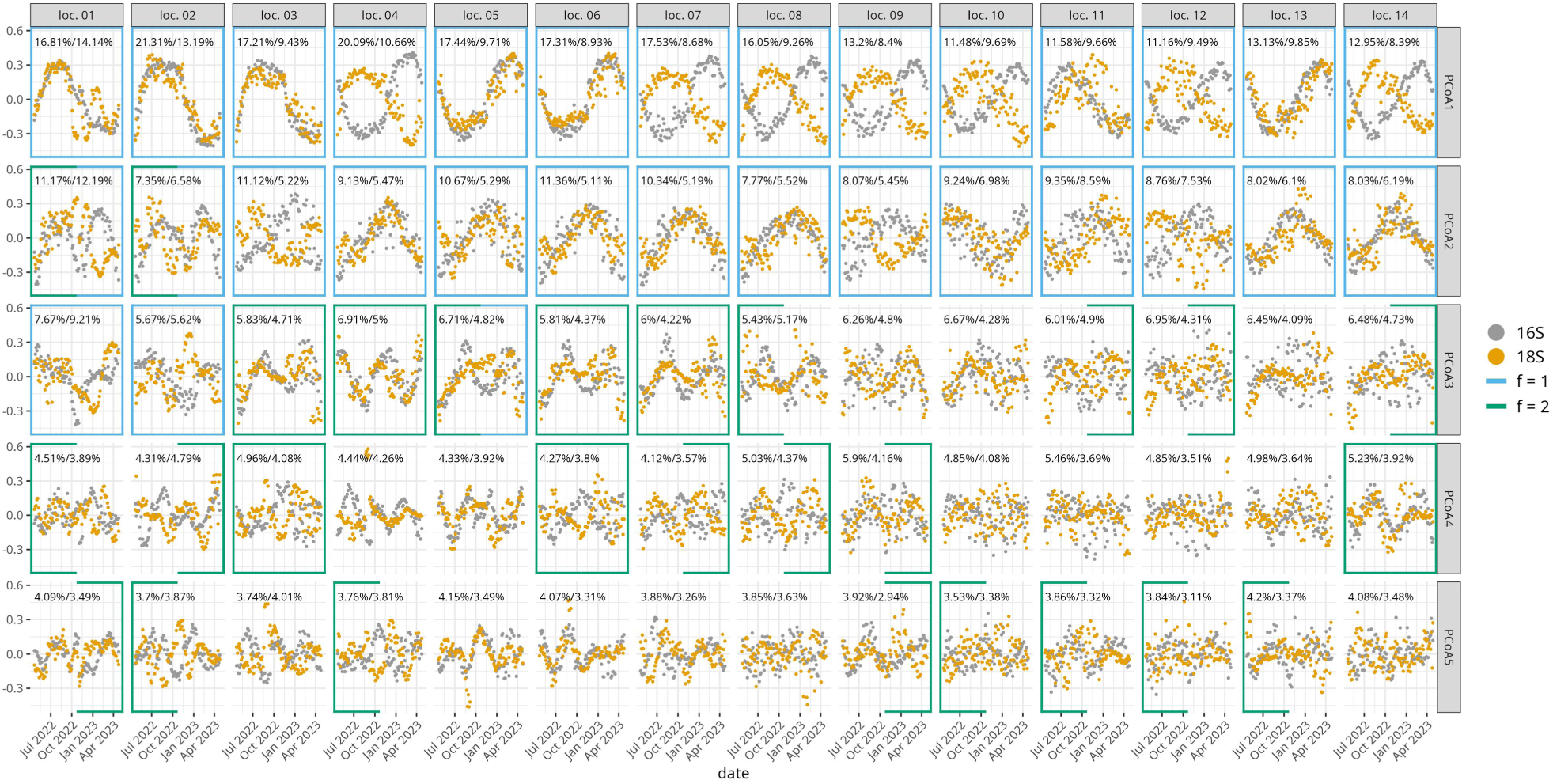
PCo axes capture sinusoidal phenologies of different frequencies across the Warnow estuary and Baltic Sea coast for both Prokaryotes (16S) and Eukaryotes (18S). Percentages give the amount of variation explained by the relevant axes for the 16S and 18S datasets, respectively. Location numbers refer to the sampling sites specified in fig. 1C. Highlighted borders around the facets indicate the identification of annual (f=1) or semi-annual (f=2) cycles using a harmonic regression in the 16S (border on the left) or in the 18S data (border on the right) presented in the facet they enclose (for details, see Methods).

To further validate the identification of the biotic SAM and to ensure that these findings are not methodological artifacts, we employed a machine-learning approach to identify the oscillations directly from community composition data. We trained Random Forests to approximate cosine waves with amplitudes of 1, frequencies of either 1 or 2, and phases regularly spaced between 0 and 360° from the biotic community composition data. The model choice was motivated by the ability of Random Forests to model non-linear relationships in high dimensional datasets and its proven track record in microbial ecology (Breiman 2001, De’ath 2007, Sperlea et al. 2021, Zschaubitz et al. 2025).

Using this approach, we detect an annual cycle at all locations but the estuarine location 6 (Fig. 3A and 3C). By contrast, the semi-annual cycle can be detected for all locations in either the 16S and the 18S rRNA gene data (Fig. 3B and 3D). The phases for which we detected the biotic SAM show a smaller spread across locations than the phases for which we detected the annual cycle (Fig 3E-H). Generally, we observe a rotation symmetry of 180° and 90° for the annual and semi-annual modes in the biotic community, respectively, which suggests that the machine learning models are insensitive to phase reversals (Fig. 3E-H). Furthermore, we detect annual and semi-annual sinusoidal patterns with a narrow range of phases, which indicates that the machine learning models discriminate between cycles with different phases. We detect the biotic SAM at similar phases but less consistently and with reduced goodness-of-fit metrics when using linear or Ridge regression models instead of Random Forests, which points to the importance of non-linear relationships in the biotic community for detecting the biotic SAM (Fig. S3).

**Fig. 3:**
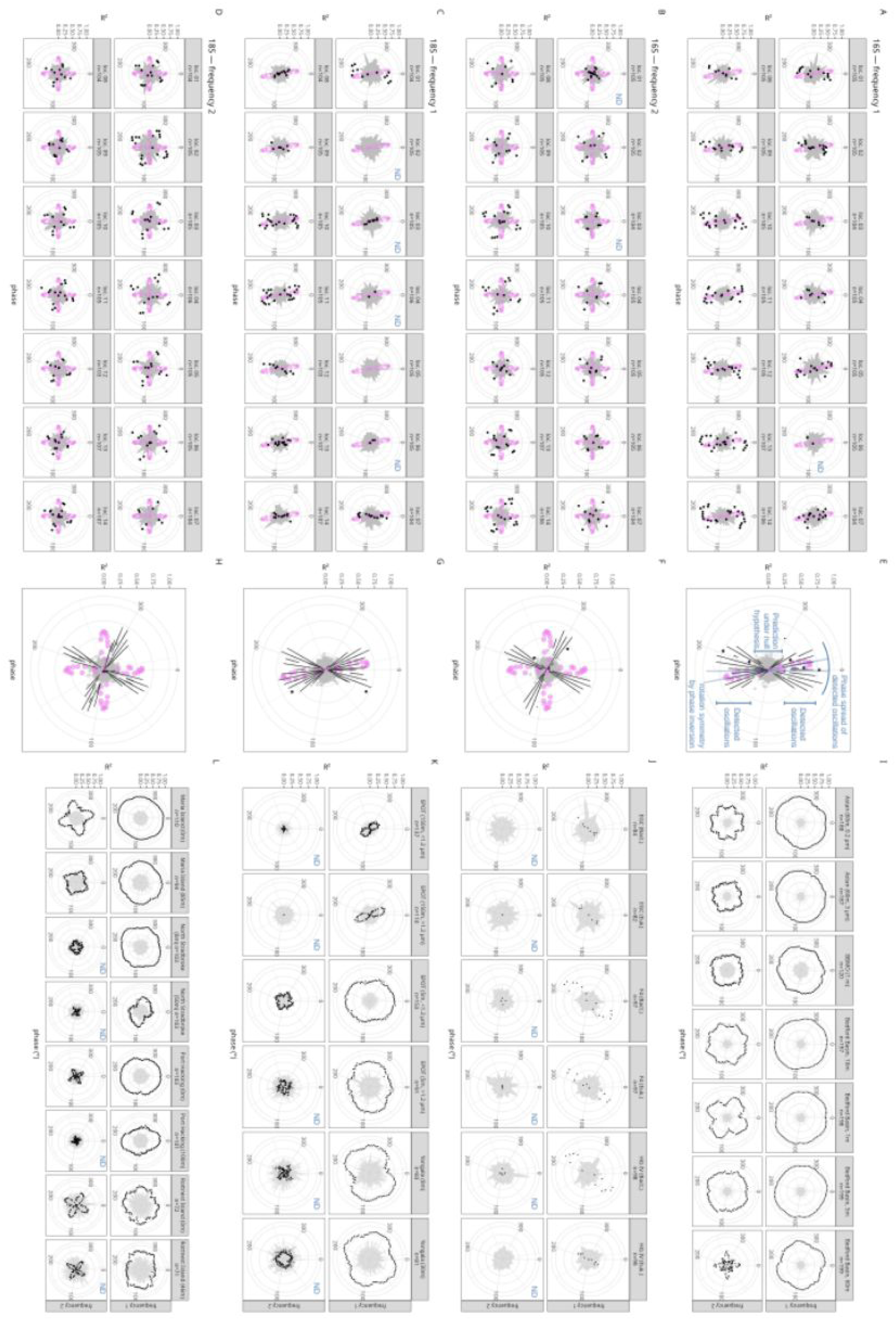
Identification of annual and semi-annual cycles in global metabarcoding datasets using a machine learning approach. Points represent the performance of a Random Forest model predicting a cosine function over time with the given frequency and phase (radial x axis) and amplitude of 1 from the Prokaryotic or Eukaryotic subset of the biotic community captured via metabarcoding (see Methods). Further description is added to **E** in blue. **A-D, I-L.** Site-specific performance of the machine learning models for the metabarcoding dataset collected in this study (**A-D**) and world-wide metabarcoding datasets (**I-L**). Black points represent models that used the biotic community composition as input. Grey areas indicate R2 values at which no oscillation would be detected, i.e., the space of R² values that are equal to or smaller than three standard deviations above the mean of the performance of control models trained on the biotic community to predict reshuffled cosine values (Methods). **E-H.** Box plots aggregate the results shown in A-D, respectively, representing Random Forest predictions of cosine values based on the biotic community composition. Grey points represent models that were trained on the biotic community composition to predict reshuffled cosine values (Methods). In **A-H**, purple points represent models that were trained on climate data. The blue annotation “ND” in the right upper corner of a facet in A-D or I-L signifies that no oscillation was detected for the setup displayed in the facet. Points with R² values below 0 are not shown.

### The biotic SAM is potentially driven by a semi-annual mode in insolation

The presence of the SAM in both Prokaryotic and Eukaryotic communities at all sampling sites along the Warnow estuary and the Baltic Sea coast, and across ecological barriers, suggests that the biotic SAM does not emerge from local interactions between organisms but is forced by external factors. Indeed, applying our machine learning approach to climate data captured at a weather station close to our sampling sites (max. 14.2 km distance, see Fig. 1C) during the year sampling was performed, we identified cycles with phases of around 350° and 0° for annual and semi-annual periods, respectively (pink dots in Fig. 3). Analysis of feature importance identified features related to either insolation or temperature for the climate SAM, and features related to insolation for the annual cycle (Fig. S4).

To verify that the climate SAM is due to insolation, we derived a formula for the dynamics of insolation from first principles (Supp. Text 1). By modelling Earth’s obliquity, rotation and revolution around the sun, this formula produces a characteristic annual pattern of insolation with annual and semi-annual modes across latitudes that is independent of the longitude of the point of interest because we average over diurnal fluctuations in insolation (Fig. 4A and B). While the annual mode of insolation has a phase reversal at the equator, its semi-annual mode has phase reversals and amplitude minima at roughly 45°N and 45°S, with a phase of 171° in the polar and a phase of −8.9° in the equatorial region (black and grey bars at frequency 1 and 2 in Fig. 4C). The model output reproduces the downward solar radiation at the nominal top of the atmosphere in the NCEP/DOE Reanalysis II to a very high degree (R²=0.94, Fig. S5). The presence of a distinct SAM in climate data as well as a semi-annual mode in insolation suggests that the biotic SAM is ultimately caused by the semi-annual mode in insolation and transmitted to the biotic communities directly as well as indirectly via climate variables such as temperature.

**Fig. 4:**
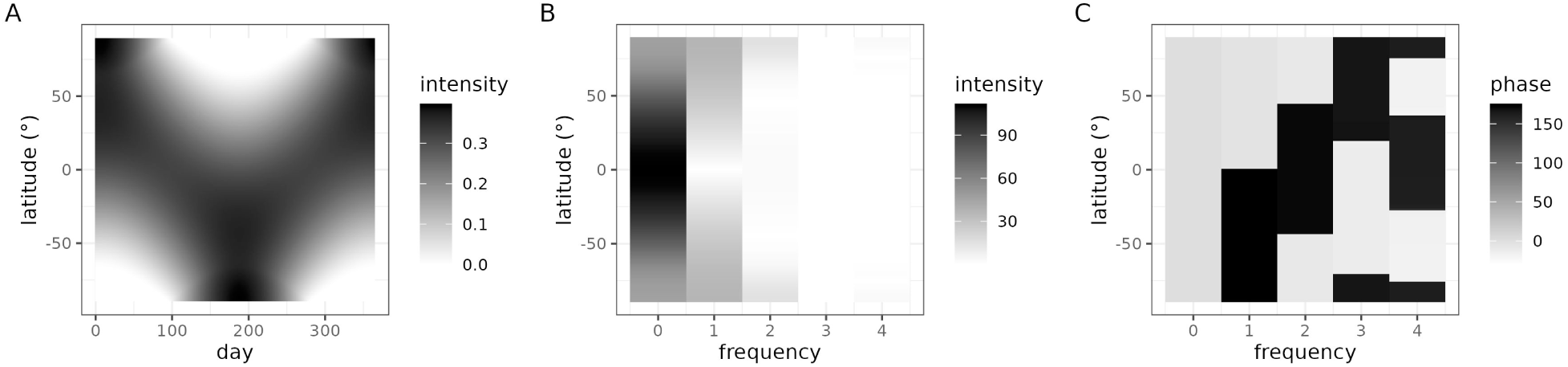
A model of insolation produces a latitudinal pattern and distinct annual and semi-annual mode. **A.** Spatiotemporal annual distribution of the intensity of the insolation derived from the insolation model (see Methods)**. B.** Amplitudes and **C.** phases of the insolation modeled using the insolation model (see Methods) derived using a numerical Fourier decomposition.

### A global survey unveils the presence of the biotic SAM in temperate zones

Owing to the role of insolation, we assume that photoautotroph organisms likely have an important role in the aetiology of the biotic SAM. Based on this and the global distribution of the semi-annual mode in insolation, we hypothesize that the biotic SAM is present globally but can be suppressed in regions where nutrient scarcity undermines photoautotrophy, i.e., in regions with a low annual average Chlorophyll a concentration (Fig. 1A). To test this hypothesis, we attempted to identify the biotic SAM using the same machine learning approach in long metabarcoding time series data collected in marine settings around the globe (Fig. 1, Tab. S1). The chosen time series datasets cover the years between 2005 and 2022, target different subgroups of the biotic community, span a latitudinal and longitudinal range of ∼120° and ∼240°, respectively, and different water column depths. To our knowledge, there are no suitable metabarcoding time series datasets available from the tropics nor the Antarctic region, nor from sediment or soil. Using this approach, we identified the annual cycle in the biotic community at all and the biotic SAM at four out of eleven time series sampling sites, all of which are in or very close to the temperate zone (white and grey dots in Fig. 1, Fig. 3I-L). Additionally, at the Port Hacking, Rottnest Island, and San Pedro Ocean Time-series (SPOT) sampling site, we detect a biotic SAM only in surface waters.

Comparing predictions of the machine learning approach between the Warnow estuary and Baltic Sea coast with the other sampling locations, we observe high predictability of the biotic SAM for only a small region in phase space for the former, whereas for locations such as SOMLIT-Astan or BBMO, cycles of any phase are predictable (Fig. 3). To test the hypothesis that this is due to the length of the time series, we truncated the time series for which we identified a biotic SAM at yearly intervals, retaining at least a full year of data. We then repeated the analysis on the truncated time series. The results show that detecting the biotic SAM requires more than a full year of sampling for most time series studied here, expect for time series with very high temporal resolution (Fig. S6).

### The biotic SAM is a community-level phenomenon

To test the hypothesis that photoautotrophs are central to the biotic SAM, we used a Pearson correlation to identify amplicon sequence variants (ASVs) in the Warnow estuary and Baltic Sea coast dataset whose phenology resembles the cycles that received the highest R² across the sampling locations for this dataset. After a Bonferroni correction and removal of all correlates with a p-value larger than 0.05, we retain 21 bacterial and 22 Eukaryote ASVs, with a maximum absolute correlation coefficient of 0.6 (Fig. 5A). Notably, 19 of the eukaryotic ASVs and 3 of the bacterial ASVs identified this way either belong to or are known to be associated with photoautotrophic genera, supporting our hypothesis. By contrast, we do not observe a direct correspondence between the biotic SAM and the intra-annual variation in Chlorophyll a concentration, contradicting a simple causal link (Fig. 5B). Furthermore, we did not identify any ASVs correlating with the SAM for location 13 for either 16S or 18S dataset or the locations 3, 5, 6, or 9 to 14 for the 18S datasets. Finally, some ASVs identified as correlates of the SAM show widely varying dynamics across locations (Fig. 5B), indicating that their abiotic and biotic environment modulates their phenology. Along similar lines, 9 out of the 30 ASVs correlating with the biotic SAM in the Maria Island time series are annotated as chloroplasts, underlining the importance of phototrophic organisms in the biotic SAM. Nevertheless, and in agreement with the results gathered for the Warnow estuary, there is no clear correspondence between these correlating ASVs, the biotic SAM, and Chlorophyll a concentrations for Maria Island (Fig S8).

**Fig. 5:**
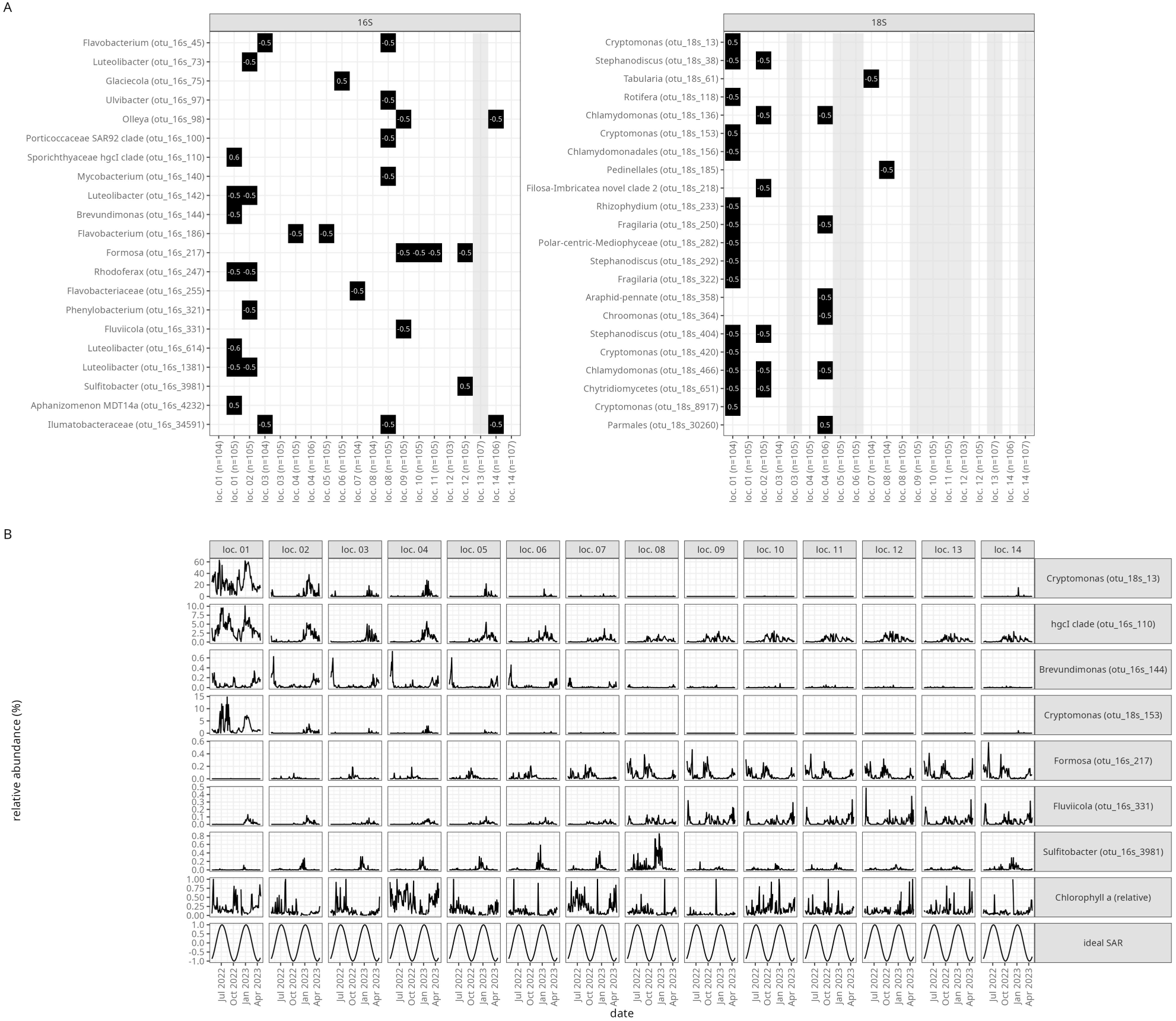
ASVs correlating with the biotic SAM do not fully explain the presence of the biotic SAM along the Warnow Estuary and Baltic Sea coast. **A.** Heatmap displaying the correlation coefficient of the relative abundance of the ASVs in question in the sampling site in question and a cosine oscillation with a phase of 27°. Only correlations significant after Bonferroni correction are displayed. Sampling sites that lack any ASV correlates are shaded in grey. **B.** Phenology of selected ASVs that correlate with the biotic SAM, relative Chlorophyll a concentration across sampling site in the Warnow Estuary dataset, and the ideal biotic SAM derived from the machine learning approach (Fig. 3). For a full list of ASVs correlating with the biotic SAM, see fig. S7.

## DISCUSSION

By combining metabarcoding sequencing time series with a newly developed machine learning approach, we report a semi-annual cycle in the biotic community in amplicon time series gathered from oceans, coastal waters, estuaries, and rivers. Due to a lack of metabarcoding time series of sufficient sampling frequency, we cannot prove the existence of the biotic SAM in non-aquatic habitats nor in the tropics.

All sampling sites at which we observe a biotic SAM are either located in the temperate zone or in the suptropics, but very close to the temperate zone (i.e., Rottnest Island, Port Hacking and SPOT). Notably, this latitudinal distribution does not reproduce the latitudinal patterns of the semi-annual mode of insolation (Fig. 1A, Fig. 4B). Because insolation is a driver of many atmospheric and climactic processes, we assume that the biotic response is not a direct one but rather relayed via, e.g., semi-annual cycles in weather variables. Furthermore, as scattering or meridional transport can induce latitudinal redistribution of solar energy in the atmosphere, the latitudinal insolation amplitude pattern observed at the exosphere cannot be expected to be conserved at the surface.

We propose that the biotic SAM is then modulated by the availability of nutrients and sunlight and that photoautotrophs play an important role in its aetiology. For one, at the SPOT sampling point, we identify the biotic SAM at the surface level but not at a water depth of 150m, which is below the euphotic zone and does not support phototrophic growth (Hickey 1992, Yeh & Fuhrman 2022). At the arctic Fram strait sampling sites, daylight is absent for around half a year, leading to an absence of photoautotroph taxa (Wietz et al. 2021, Priest et al. 2025). At the Australian East Coast, we attribute the absence of the biotic SAM at the three northern sampling locations and the presence of the biotic SAM at the southernmost one to the interplay of oligotrophic and nutrient-rich water currents. That is to say, the Yongala and North Stradbroke Island sampling locations are fed by the East Australian Current and its extension, which is an arm of the Chlorophyll a-depleted Atlantic subtropical gyre (Fig. 1A, Fig. 1D) (Lynch et al. 2014, Oliver et al. 2016, Brown et al. 2018). By contrast, the water at the Maria Island sampling station, at which we identify the biotic SAM, is dominated by the Zeehan current that is not part of the subtropical gyre. The Port Hacking sampling station is located at the border of the nutrient-deficient area generated by the subtropical gyre and displays the biotic SAM at the sea surface. At both the SPOT and Maria Island sampling stations, surface water algal blooms are linked to upwelling events (Ajani et al. 2001, Thompson et al. 2009, Cram et al 2015). Furthermore, the temperate zones have a higher phytoplankton species turnover than the subtropics or tropics, allowing for more elaborate ecological dynamics (Righetti et al. 2019). We speculate that there is a close relationship between the biotic SAM described here and the dynamics of phytoplankton blooms that can occur twice in a year (Winder & Cloern 2010, Reinl et al. 2023, Zhang et al. 2017).

The variability observed in the timing of blooms suggests that, while the biotic SAM may set the window of opportunity for a bloom, the latter is only realized if further preconditions are met. The biotic SAM in the Warnow estuary has extreme (i.e., maximal or minimal in contrast to close-to-zero) values in July and January and October and April. In the Baltic Sea, diatom blooms take place, on average, between March and May with a second, lower abundance peak around October, whereas cyanobacterial blooms last from June to August (Snoeijs et al. 2016, Angeler et al. 2019, Beltran-Perez & Waniek 2021). Therefore, the biotic SAM might be used to set expectations regarding the onset of algal blooms, making it possible to see that an algal bloom failed to materialize and ask why it did so. Similarly, the biotic SAM might be instrumental in describing the succession of blooms of heterotrophic bacteria that relays the energy and nutrients in algal polysaccharides through the biotic community (Teeling et al. 2012, 2016, Bunse & Pinhassi 2017). Furthermore, we observe a certain degree of variation in the phases identified for the biotic SAM in the truncated time series analyzed, pointing to temporal variation in the biotic SAM (Fig. S6). The site with the lowest amount of variation is the BBMO, for which the phase of the SAM only switches between ∼25° (for time series ending in the years 2007, 2008, 2011, and 2012) and ∼65° (for time series ending in the years 2009 and 2010). This stability echoes the high inter-annual stability of the seasonal biological patterns at the BBMO site (Vallina et al. 2023), underlining the potential use of the biotic SAM in biomonitoring.

While our analyses suggest that photoautotrophs are central to the aetiology of this biotic SAM, it cannot be explained on the level of single ASVs as there are sampling locations for which we do not identify any ASVs correlating with the biotic SAM (Fig. 5). Instead, implicitly capturing interactions between ASVs are essential to detecting the biotic SAM. This is supported by our inability to detect the biotic SAM in the Bray-Curtis distances alone and our ability to do so after a PCoA (Figs., 2, S1, S2), as well as our inability to detect the biotic SAM using linear machine learning models in contrast to Random Forest models (Fig. 3, S3).

Based on these methodological results, we argue that the biotic SAM is a community-level dynamic that cannot be explained by single taxa. That is to say, biotic communities in aquatic environments are exposed to the semi-annual mode in insolation as an environmental signal. The response of the local biotic community to this signal depends on its composition, for example on the proportion of photoautotrophs present, leading to specific and divergent manifestations of the biotic SAM in different locations, even spatially very close ones. For example, we observed that the phenologies of some ASVs that correlate with the biotic SAM are different at other sampling locations, likely owing to differences in local abiotic factors (e.g., differences in salinity and hydrodynamics) and the presence of interacting species (Fig. 5). Thus, our results suggest that the biotic SAM is not emergent from biotic interactions as previously proposed (Caracciolo et al. 2022), but rather owes to the interplay between an external cause (i.e., insolation) and non-linear biotic interactions within the community. This proposed causal link between the semi-annual mode in insolation and the biotic SAM makes it possible to use the latter to monitor changes affecting ecosystems across scales, from the local to the planetary: Any variation in the presence or phase of the biotic SAM that cannot be explained by insolation models derived from first principles will be due to changes in atmospheric composition, climatological processes or differences between water currents in stratified water bodies. However, this kind of analysis requires metabarcoding time series with a high sampling frequency, which are currently rare.

## CONCLUSION

To conclude, our analyses suggest that the biotic SAM is a hitherto unrecognized phenomenon in aquatic biotic communities that links phenomena across scales, from the planetary to the microbial. We strongly suppose that it is a community-level response to insolation modulated by the availability of nutrients. This finding serves as a reminder that ecological dynamics observed at the local level can be due to properties of the whole Earth system. As such, the biotic SAM might enable us to capture environmental change by proxy, using the dynamics of biotic communities to monitor the dynamics of the whole Earth system on a local scale.

## Author contribution statement

TS and ML designed the study; TS analysed and visualized the data; TS and ML wrote the initial draft of the manuscript; CCG, LV, EZ, and CN performed the sampling and the measurements; SK derived the insolation model; SE, CL, and CH performed the metabarcoding analyses; HS provided the data storage infrastructure and developed the machine learning approach with TS; JD provided an explanation for the global distribution of the biotic SAM in the data; ML supervised the study. All authors participated in reviewing and editing the manuscript.

## Data accessibility statement

Metabarcoding data from the Warnemünde estuary and Baltic Sea coast are publicly available within the European Nucleotide Archive (ENA) under accession IDs PRJEB88008 (16S metabarcoding data) and PRJEB88011 (18S metabarcoding data), and PRJEB88048 (umbrella project). Chlorophyll a data is publicly stored in the IOWDB and can be accessed via https://doi.iow.de/10.12754/data-2025-0001. Code for sequence procession is available under https://git.iow.de/bio_inf/IOWseq000042_OTCgenomics_seqs. R and Python code used for analyses and to create the figures displayed in this study is available under https://dx.doi.org/10.6084/m9.figshare.27249597.

Climate data from the weather station Rostock-Warnemünde was downloaded from the Deutscher Wetterdienst (DWD) Climate Data Center Open Data ftp server (https://opendata.dwd.de/climate_environment/CDC/) as daily averaged values on November 30th, 2023. Data points with missing data (indicated by a value of −999) were removed from the analysis. Chlorophyll a concentration data for the year 2021 were downloaded from https://zenodo.org/records/7092220 (Yu et al. 2023). Processed data on the microbial community composition in the form of denoised ASVs was downloaded from the Australian Microbiome Initiative page in the Bioplatforms Australia Data Portal (https://data.bioplatforms.com/organization/about/australian-microbiome) on Jan. 26th, 2024 (Brown et al. 2018). Only marine samples from the sampling stations Maria Island, Port Hacking, North Stradbroke Island, Rottnest Island, and Yongala were used. The same rarity filter that was used for the Warnow estuary data set was also applied here. ASV tables for the SOMLIT-Astan (Caracciolo et al. 2022) and BBMO (Giner et al. 2019, Krabberød et al. 2022, Vallina et al. 2023) time series were downloaded from dx.doi.org/10.5281/zenodo.5032450 and https://github.com/ramalok/BBMO.krabberod.etal/tree/main, respectively. Raw data for these are publicly available at the European Nucleotide Archive (ENA) under the accession numbers PRJEB48571, PRJEB23788 and PRJEB38773. Raw reads for the SPOT dataset (Yeh & Fuhrmann 2022) were downloaded from ENA via the accession numbers PRJEB48162 and PRJEB35673 and processed as described for the Warnow estuary data set. Processed ASV tables for the Fram strait sampling sites F4, EGC and HG-IV (Priest et al. 2025, Wietz et al. 2021) and for the Bedford Basin sampling sites (Raes et al. 2022) were downloaded from https://github.com/matthiaswietz/FRAM_eDNA and https://github.com/EricRaes/Bedford_Basin_Time_Series, respectively. Raw sequences for these are available at the ENA under the accession numbers PRJEB43905 and PRJNA785606, respectively.

Shapefiles for Mecklenburg-Vorpommern were downloaded from http://download.geofabrik.de/europe/germany.html; for visualization purposes, we used the shapefiles for land use, bodies of water and natural features.

## ACKNOWLEDGEMENTS

We want to thank Martin Schmidt, Alexander Tagg and Stefan Lüdtke for fruitful discussions. We want to thank David Riedinger, Mariano Santoro, Lara Renner, Julian Lino Peter Kunis, Finn Spieß, Anne Lea Schenk, Lisa Gaertner, and Heike Benterbusch-Brockmöller for their support in sampling.

We would like to acknowledge the contribution of the Australian Microbiome consortium in the generation of data used in this publication. The Australian Microbiome initiative is supported by funding from Bioplatforms Australia and the Integrated Marine Observing System (IMOS) through the Australian Government’s National Collaborative Research Infrastructure Strategy (NCRIS), Parks Australia through the Bush Blitz program funded by the Australian Government and BHP, and the CSIRO.

