## Supplemental Information for "Semi-Annual Cycles in the Biotic Communities of Temperate Aquatic Habitats"

### Supplementary Information for "Semi-Annual Cycles in the Biotic Communities of Temperate Aquatic Habitats."

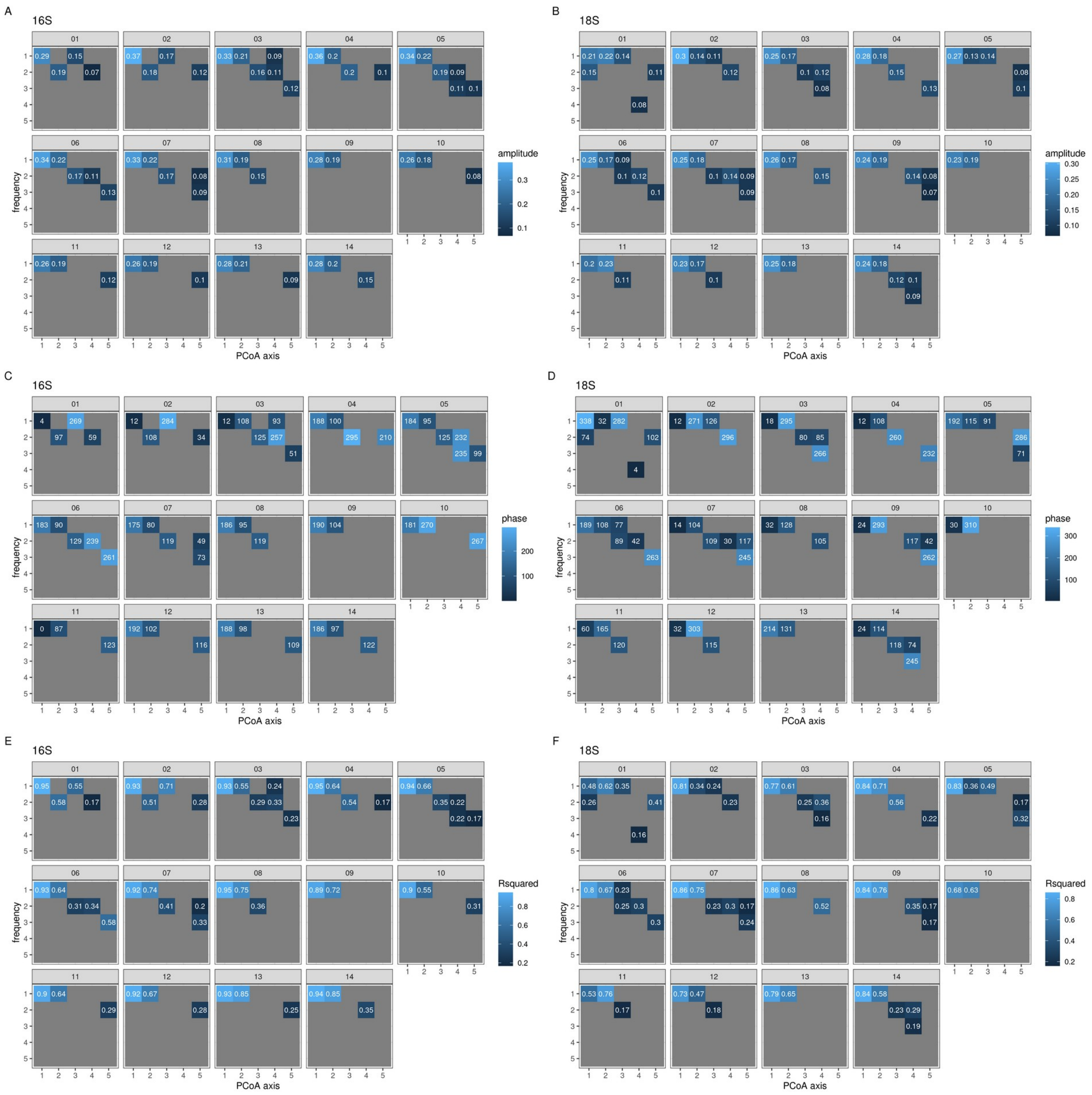


**Fig. S1.**

Harmonic Analysis on Bray-Curtis derived PCoAs of the metabarcoding data from the Warnow estuary and Baltic Sea coast validates the presence of a biotic SAM. Amplitudes (A, B), phases (C, D), and $R^2$ compared to the original PCoA data (E, F) are shown for those combination of PCo axis and frequency that have a z-score of more than 10 (see methods).

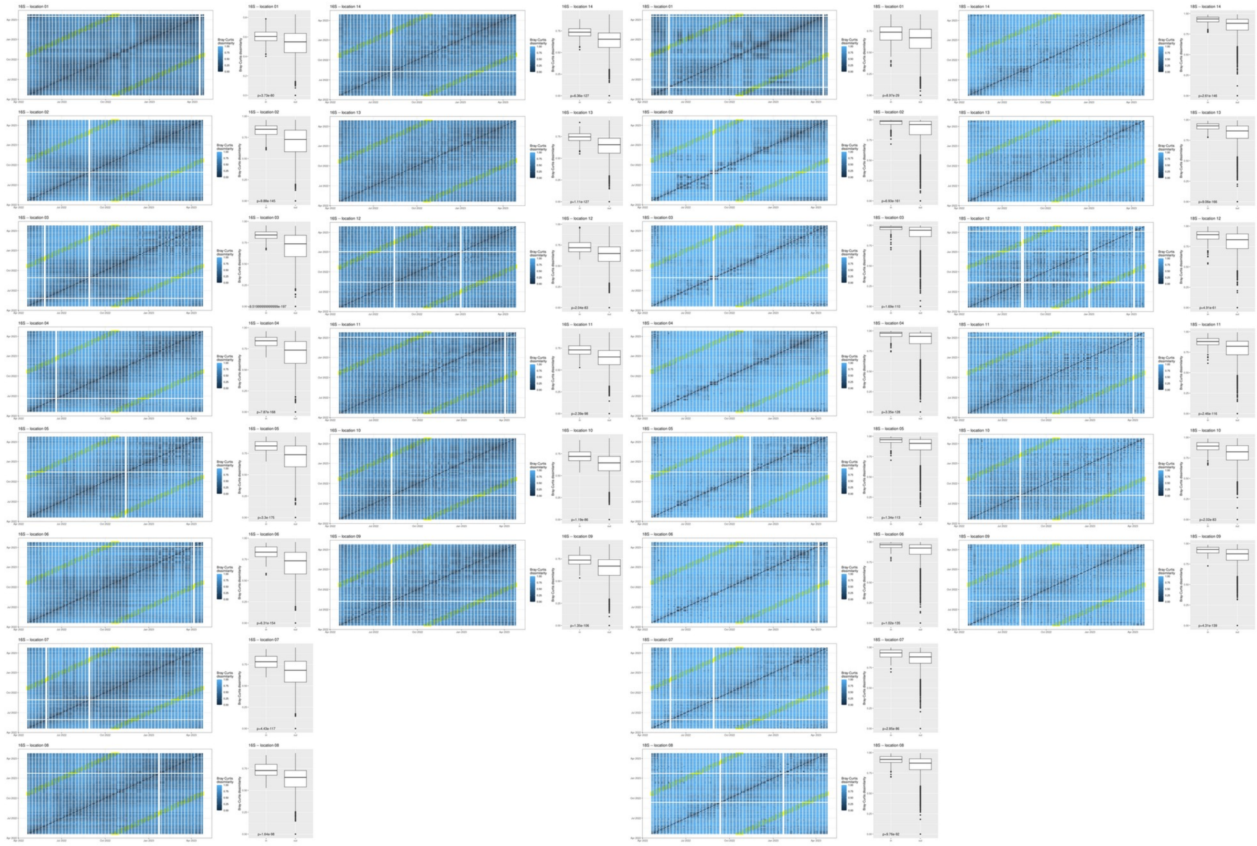

**Fig. S2.**

A simple statistical approach cannot detect SAMs on Bray-Curtis matrices directly. Heatmaps show Bray-Curtis matrices, separated by metabarcoding target gene and sampling location, sorted by date; boxplots show the distribution of Bray-Curtis dissimilarities among samples separated by half a year (highlighted in yellow in the heatmaps; category “in”) and all other pairs of samples (category “out”). Consistently, these two groups’ means are different according to a t-test; however, the mean Bray-Curtis dissimilarity of the “in” group is consistently higher than the mean Bray-Curtis dissimilarity of the “out” group.

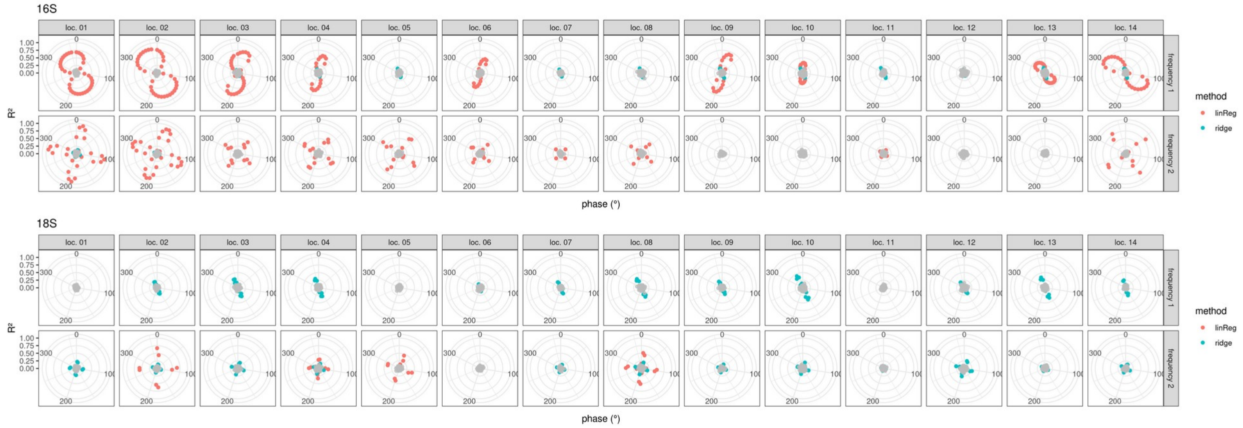


**Fig. S3.**

Linear models detect biotic annual and semi-annual cycles at fewer locations along the Warnow estuary and the Baltic Sea coast. Data were generated as for fig. 3 but using the models LinReg (red dots) and ridge (blue dots) from scikit-learn instead of a Random Forest model. Grey dots represent controls generated by resampling the cosine target.

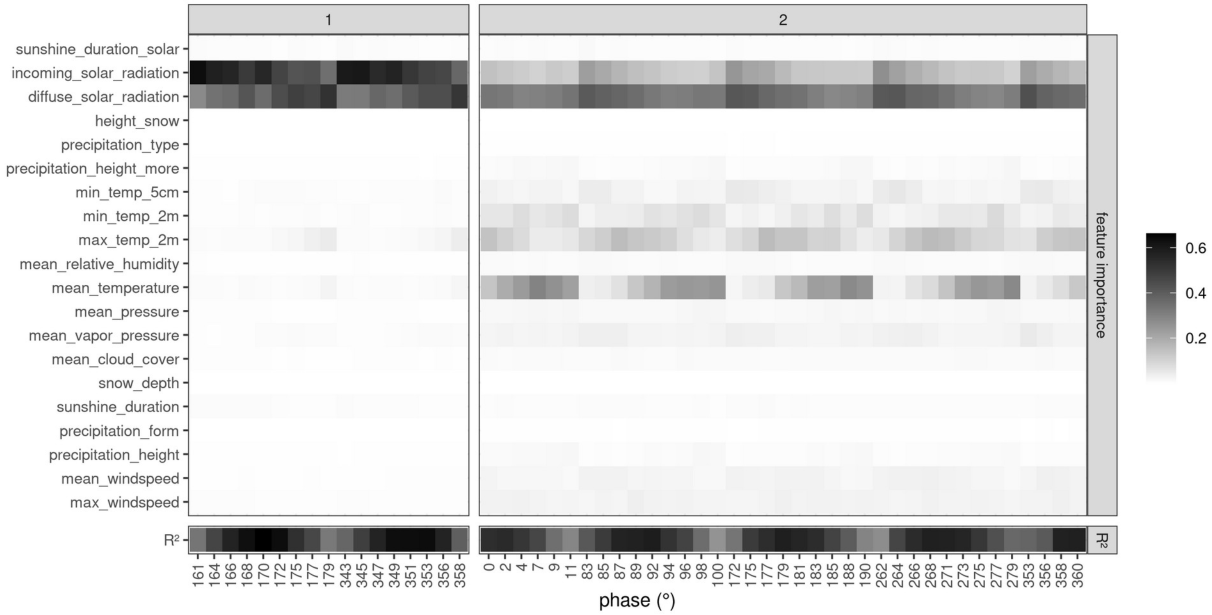


**Fig. S4.**

A feature importance analysis for the cycles in climate data points to solar radiation and temperature. Feature importance analysis shown for all models with $R^2>0$.

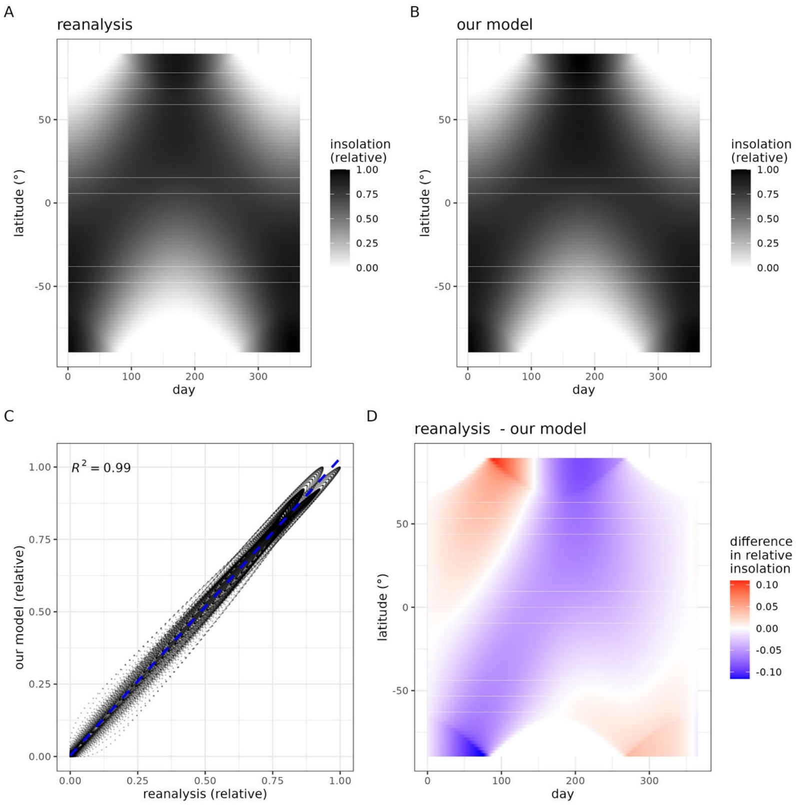


**Fig. S5.**

The insolation model proposed in this paper almost perfectly reproduces the downward solar radiation at the nominal top of the atmosphere (dswrf.ntat) data present in the NCEP/DOE Reanalysis II. A and B. Latitudinal and temporal distribution of insolation relative to the maximal value for the Reanalysis (A) and our model (B). C. Direct composition of the data presented in A and B. D. Latitudinal and temporal distribution of the error between the values presented in A and B.

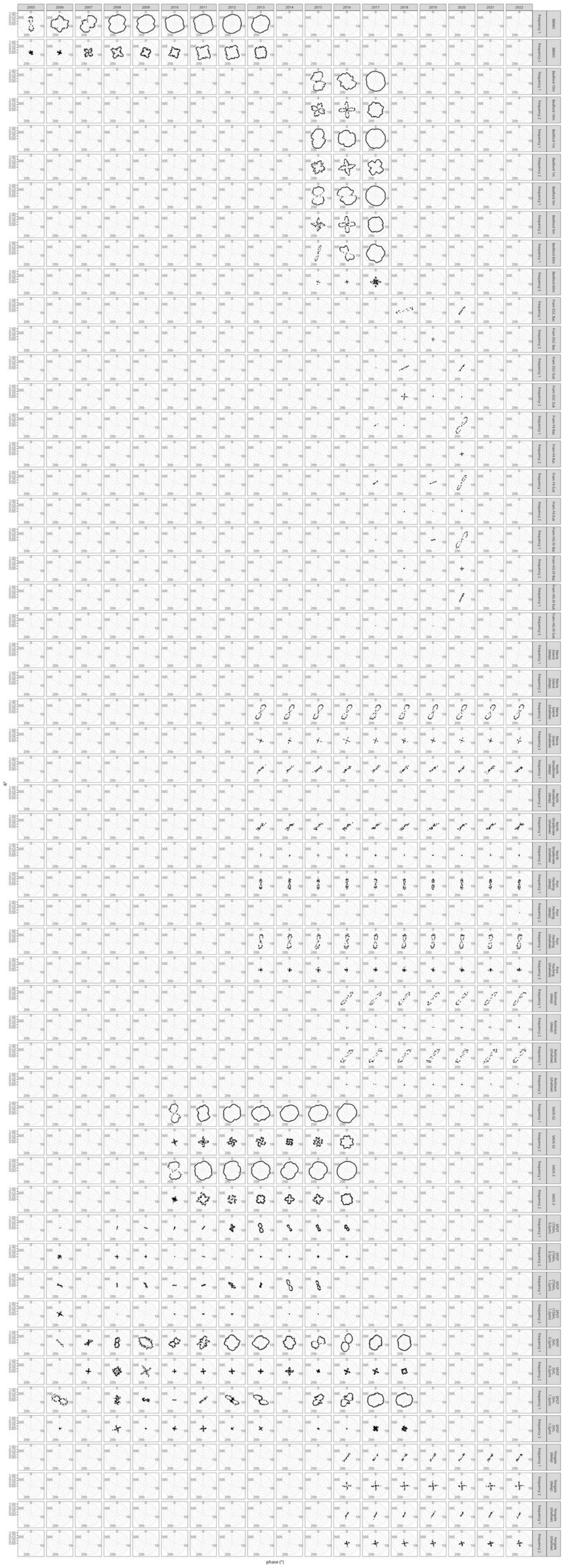

**Fig. S6.**

Results of the machine learning approach for identifying annual and semi-annual cycles for time series truncated after each year of data. Black dots represent model performances; only results with $R^2>0$ are shown.

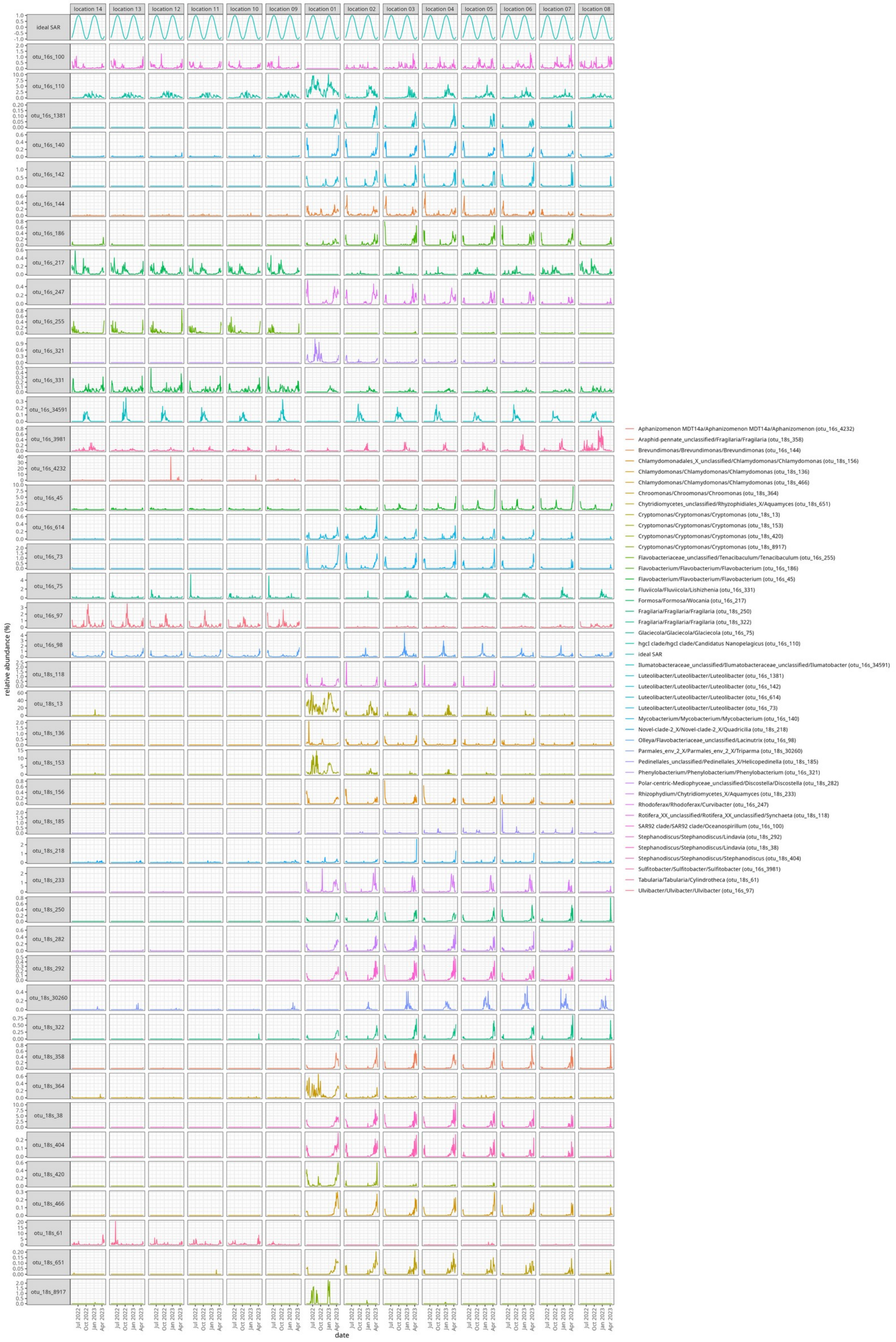


**Fig. S7.**

Phenology of all ASVs correlating with the optimally predictable biotic SAM the Warnow estuary and the Baltic Sea coast.

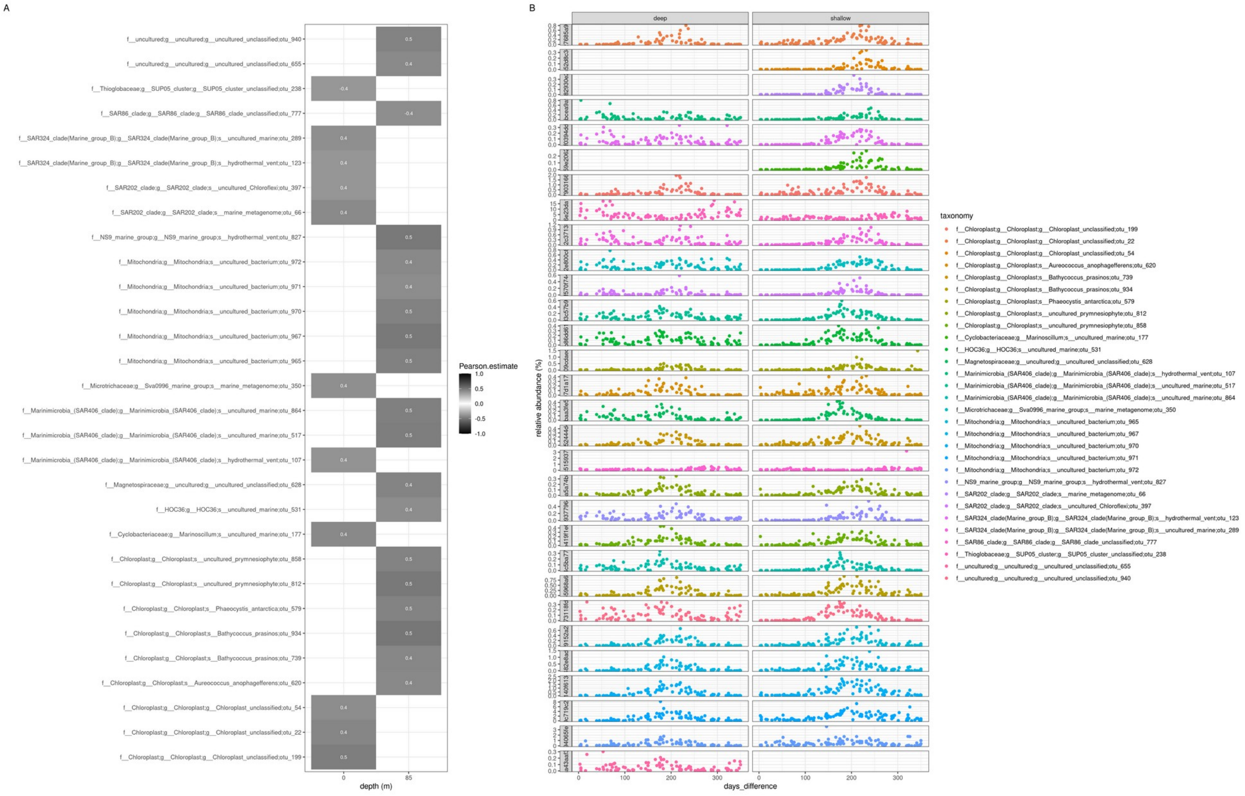


**Fig. S8.**

Correlation coefficients and phenology of ASVs correlating with the optimally predictable biotic SAM for the Maria Island time series. A. Heatmap displaying the correlation coefficient of the relative abundance of the ASVs at the sampling depth in question and a cosine oscillation with a phase of 27°. Only correlations significant after Bonferroni correction are displayed. B. Phenology of all ASVs that correlate with the biotic SAM.

**Table S1.** Information on the metabarcoding datasets used in this study.

| Name | Location | Time range & frequency | Depth | Target gene | Size fraction | References |
|---|---|---|---|---|---|---|
| BBMO | Northwestern Mediterranean Sea (41º 40' N, 2º 48' E) | 2004-2013, monthly | ~1 m | V4 of 16S rDNA & V4 of 18S rDNA (combined) | 0.2–3 µm & 3–20 µm (combined) | (25, 50*, 51*) |
| Bedford Basin | Estuary to the North Atlantic (44° 41' N 63° 38' W) | 2014-2017, weekly | 1 m, 5 m, 10 m, 60 m | V4-V5 of 16S rDNA | >0.2 µm | (*52*) |
| Fram Strait EGC, F4, HG-IV | Greenland Sea (79° 00 N, 5° 24' W; 79° 00 N, 7° 00 E, 79° 01 N 4° 15' E) | 2016-2020, monthly | 65-150 m, 20-110 m, 24 m – 131 m | V4–V5 of 16S rRNA, V4–V5 of 18S | >0.2 µm | *(10, 28)* |
| Maria Island | 42° 35' S, 148° 14' E | 2012-2022, monthly | 0 m, 10 m, 20 m, 50 m, 75 m, 85 m | V1-3 for 16S (Bacteria and Archaea), V4 of 18S r DNA (combined) | >0.2 µm | (*31*) |
| North Stradbroke Island | 27° 20' S, 153° 33' E | 2012-2022, monthly | 0 m, 50 m | See Maria Island | >0.2 µm | (*31*) |
| Port Hacking | 34° 05' S, 151° 15' E | 2012-2022, monthly | 0 m, 100 m | See Maria Island | >0.2 µm | (*31*) |
| Rottnest Island | 32° 00' S, 115° 25' E | 2015- 2022, monthly | 0 m, 46 m | See Maria Island | >0.2 µm | (*31*) |
| SPOT | Pacific Coast (33° 33' N, 118° 24' W) | 2005-2018, monthly | 5 m or 150 m | V4-V5 of 16S rDNA and V4-V5 of V18S rDNA (combined) | 0.2–1 or 1–80 µm | (*27*) |
| SOMLIT-Astan | English Channel (48°46′18″ N, 3°58′6″ W) | 2009-2016, bimonthly | 60 m | V4 of 18S rDNA | 0.2–3 µm or 3–20 µm | (*23*) |
| Warnow Estuary | 54°05' - 54°08' N, 11°52' - 12°09' S | Apr. 2022-Apr. 2023, Twice a week | 0 m | V4 of 16S or V4 of 18S | >0.2 µm | (*20*) |
| Yongala | 19° 18' S, 147° 37' E | 2015-2022, monthly | 0 m, 30 m | See Maria Island | >0.2 µm | (*31*) |
